# Complementary topology of maintenance and manipulation brain networks in working memory

**DOI:** 10.1101/371294

**Authors:** S.W. Davis, C.A. Crowell, L. Beynel, L. Deng, D. Lakhlani, S.A. Hilbig, W. Lim, H. Palmer, D. Nguyen, A. V. Peterchev, B. Luber, S.H. Lisanby, L.G. Appelbaum, R. Cabeza

**Affiliations:** Department of Neurology, Duke University School of Medicine, Durham, NC; Department of Psychiatry and Behavioral Science, Duke University School of Medicine, Durham, NC; Center for Cognitive Neuroscience, Duke University, Durham, NC; Department of Biomedical Engineering, Duke University, Durham, NC; Department of Electrical and Computer Engineering, Duke University, Durham, NC; Department of Neurosurgery, Duke University School of Medicine, Durham, NC; Department of Psychology & Neuroscience, Duke University, Durham, NC; National Institute of Mental Health, Bethesda, MD

**Keywords:** Working Memory, Manipulation, Maintenance, Network Connectivity, fMRI

## Abstract

Working memory (WM) is assumed to consist of a process that sustains memory representations in an active state (*maintenance*) and a process that operates on these activated representations (*manipulation*). Prior fMRI studies have examined maintenance and manipulation in separate task conditions, whereas in real life these processes operate simultaneously. In the current study, the neural mechanisms of maintenance and manipulation were disentangled during the same task by parametrically varying these processes. During fMRI, participants maintained consonant letters in WM while sorting them in alphabetical order. Maintenance was investigated by varying the number of letters held in WM and manipulation by varying the number of moves required to sort the list alphabetically. The study yielded three main findings. First, the degree of both maintenance and manipulation demand had significant effects on behavior that were associated with different cortical regions: maintenance was associated with bilateral prefrontal and left parietal cortex, and manipulation with right parietal activity, a link that is consistent with the role of parietal cortex in symbolic computations. Second, univariate fMRI and tractography based on diffusion-weighted imaging showed that maintenance and manipulation regions are supported by two dissociable structural networks. Finally, maintenance and manipulation functional networks became increasingly segregated with increasing demand, possibly reflecting the protection of information held in WM from interference generated by manipulation operations. These results represent a novel approach to study the brain as an adaptive system that coordinates multiple ongoing cognitive processes.

**Significance Statement:** Despite the importance of working memory (WM) in everyday life, little is known about how the brain is able to simultaneously maintain and manipulate information stored in short-term memory buffers. We examined evidence for two distinct, concurrent cognitive functions supporting maintenance and manipulation abilities by testing brain activity as participants performed a WM alphabetization task. We found behavioral and neural evidence in support of dissociable cognitive functions associated with these two operations. Furthermore, we found that connectivity between these networks was increasingly segregated as difficulty increased, and that this effect was positively related to individual WM ability. These results provide evidence that network segregation may act as a protective mechanism to enable successful performance under increasing WM demand.

## Introduction

Working memory (WM) is a fundamental ability that allows humans to process information not currently available to the senses. WM is typically assumed to consist of dissociable *maintenance processes*, which sustain information in an activated state, and *manipulation processes,* which operate on the maintained information (Baddeley, 2000). This distinction is fundamental for WM theory and it has been the focus of multiple fMRI studies (D’Esposito et al., 1999a; Postle et al., 2006; Rypma, 2006). However, most fMRI studies comparing maintenance and manipulation operations have been limited in three ways. First, maintenance and manipulation have usually been investigated in separate tasks or conditions (Rypma et al., 1999; Postle et al., 2006), while in real life, they occur simultaneously. Second, the specific computations underlying manipulation have rarely been operationalized or examined. For example, in the typical manipulation task in which participants are asked to put letters in alphabetical order (D’Esposito et al., 1999b; Bunge et al., 2000), the critical operation is mentally shifting the position of each letter into a new order. The difficulty of this process depends on the number of “sorting steps” needed to achieve the reordering, which is a factor that has not been investigated in behavioral or fMRI studies of WM. Finally, most WM maintenance-manipulation fMRI studies have focused on univariate activity and have not examined functional interactions among multiple regions. Given that WM requires rapid exchange of information among many regions, characterizing the connectivity patterns between these systems is essential for understanding the processes that enable maintenance and manipulation of information in WM.

The current study addressed these three problems. To address the first two interrelated limitations, this study investigated maintenance and manipulation during the same *Delayed Response Alphabetization Task* (DRAT), which utilizes both forms of WM processing. Here maintenance was examined by assessing parametric changes in the number of letters held in WM (*Set Size*) and manipulation, by assessing the number discrete moves required to alphabetize the letters (*Sorting Steps*), both during the delay period. It was hypothesized that Set Size and Sorting Steps would have distinct effects on performance and elicit distinct parametric patterns of univariate activity. Based on neuroimaging evidence linking Set Size to prefrontal cortex (PFC; for review, see Rypma and D’Esposito, 1999), and abstract symbol manipulations to the superior parietal lobule (SPL; Postle et al., 2006), a dissociation between these two regions was expected.

Lastly, to address the limited focus of previous maintenance-manipulation fMRI studies on univariate activity, we also examined network dynamics. Graph measures of network segregation and reconfiguration (D’Esposito et al., 1999b; Han and Kim, 2004; Eriksson et al., 2015) were used to describe the dynamics of maintenance and manipulation networks as a function of maintenance or manipulation demands. Changes in the relational complexity of a task have been associated with variations in the segregation of PFC regions (Harvey et al., 2013; Cohen and D’Esposito, 2016), as well as to more global alterations in the organization of whole-brain partitions (Chan et al., 2014; Cohen et al., 2014). The present study offers an intermediate approach between these local and global scales, defining widespread, task-related networks that represent concurrent maintenance and manipulation operations. Given that the goal of maintenance is to sustain information in the same state whereas the goal of manipulation is to alter this state, it was expected that negative association would exist between networks supporting these processes. Moreover, it was also expected that this segregation of processing would increase with task difficulty.

In sum, we hypothesized that Set Size and Sorting Steps would (1) have differential effects on WM performance, (2) be associated with univariate activations in different brain regions (e.g., PFC vs. SPL), and (3) be supported by dissociable neural networks. We expected that the answer to these hypotheses would clarify the neural mechanisms underlying the two main types of cognitive operations mediating working memory function, maintenance and manipulation.

## Materials and Methods

### Participants

Forty-four young adults aged 18 to 35 (mean 22.8 ± 4.6, 23 F) participated in the study for monetary compensation and consented to the protocol approved by the Duke Medical School IRB. Participants had no history of psychiatric or neurological disorders and were not using psychoactive drugs. These participants were enrolled in a 6-day TMS protocol, but only data from the Screening session (Day 1) and MR Imaging (Day 2) are reported here. Three individuals were excluded because of poor functional imaging quality (due to excessive movement or falling asleep during the scan), and hence 41 participants are included in the analyses.

### Behavioral procedures

The study investigated a *Delayed Response Alphabetization Task* (**Figure 1**). In this task, an array consisting of 3-9 consonant letters was presented for 3 seconds followed by a 5-second delay period during which participants mentally reorganized letters into alphabetical order. Vowels were excluded to prevent chunking. After the delay period, a letter and number were presented together for 4 seconds and the participants pressed one of three buttons to indicate if the probe letter (1) appeared in the position indicated by the number in the alphabetized list (*Valid*, 40% of trials), (2) was part of original set but the number did not match the position in the alphabetized list (*Invalid*, 40% of trials), or (3) was not part of the original set (*New*, 20% of trials). These three types of trials occurred in random order. For all three conditions, the probe was never from the first half of the alphabetized array, and in the Invalid condition, to exclude obvious differences between correct and incorrect position, the number above the letter was always within 1 step of the letter’s actual alphabetized position. During the subject-specific titration on Day 1 (see the following paragraph for more information), the response phase was followed by a 5-second (mean) inter-trial interval (ITI). During practice (10 trials), participants were given feedback during this ITI on the accuracy of their previous trial response. Twenty-five trials were included in each of the 6 blocks with a brief, self-paced rest interval between blocks.

**Figure 1.**
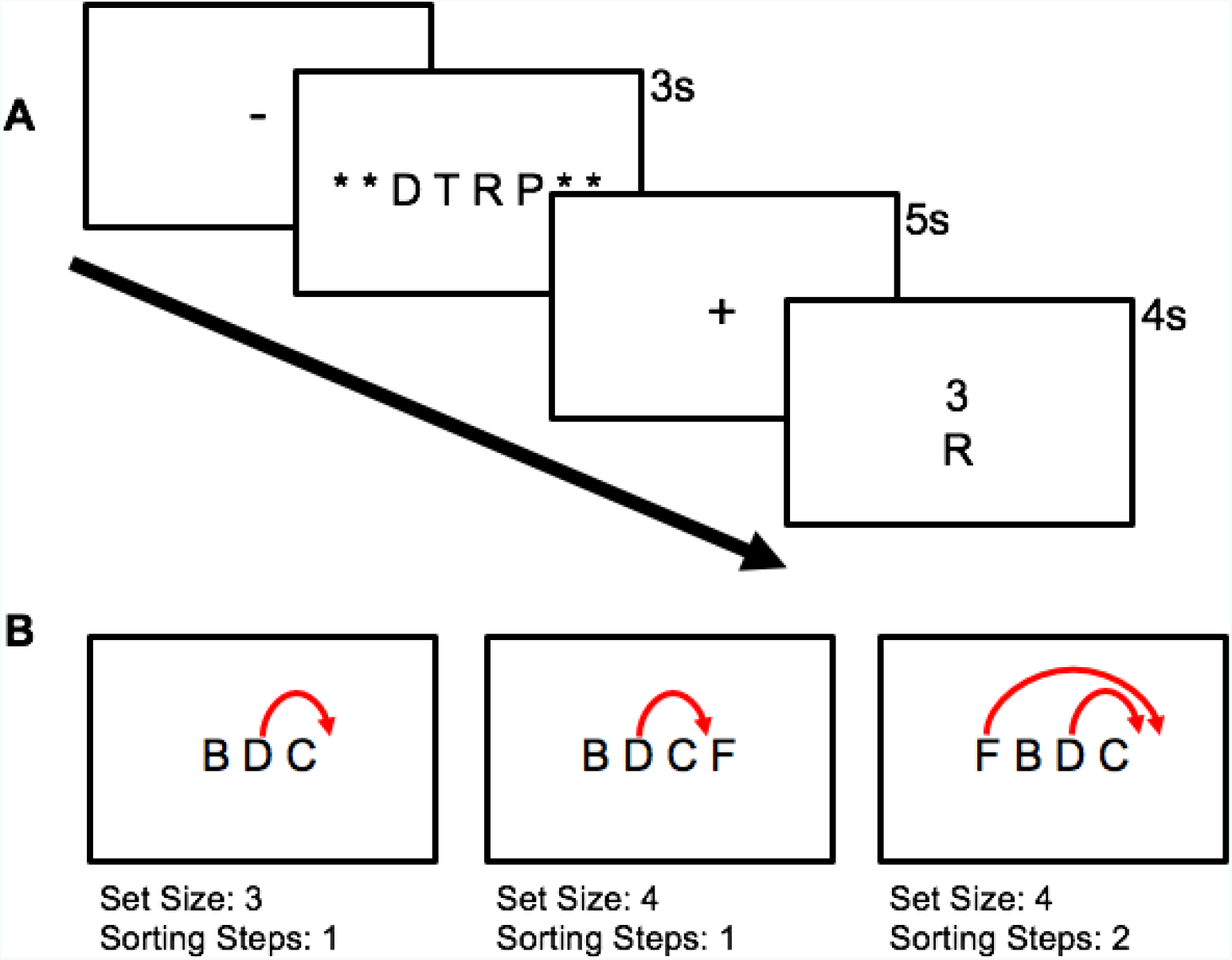
Illustration of Delayed Response Alphabetization Task with (**A**) stimulus sequence and (**B**) and a schematic illustration of the variation in the minimum number of necessary sorting steps across two different Set Sizes. Notice that while the 2nd and 3rd trials have an equal number of letters, the minimum number of steps necessary to alphabetize the array increases from 1 to 2.

As part of the overall protocol, subjects participated in 6 experimental sessions, but only the first two are relevant to this study. In the first session, participants performed the DRAT outside the scanner, while seated at a computer terminal, in order to identify the range of Set Size optimal to each participant. The optimal Set Size was identified using 2-down-1-up staircase procedure: when a trial was answered correctly, the Set Size was increased by 1, and when it was answered incorrectly, the Set Size was decreased by 2. Accuracy data for each Set Size was then fitted to a sigmoid function, with Criterion set at 82% accuracy. The two Set Sizes with sigmoid-fitted accuracy immediately greater than Criterion were defined as *Very Easy* and *Easy* levels, and the two Set Sizes with accuracy below Criterion were defined as *Medium* and *Hard* levels. Thus, the four Set Size levels selected for an individual depended on his/her WM ability (e.g., 3-4-5-6 letters in one participant, 4-5-6-7 in another participant). This method balanced task demands across participants. To ensure that the psychometric function was not strongly influenced by noise for Set Sizes with a low number of trials, 50% accuracy was used for the largest set sizes if less than 10 trials were tested. To achieve more stable curve fits, peripheral anchors were added by including points for Set Sizes of 1 and 2 at 100% accuracy and Set Sizes 10 and 11 at 50% accuracy.

In the second session, participants performed the DRAT inside the scanner. Four blocks, each with 30 trials, were performed using the 4 difficulty levels defined from session 1 performance, with equal numbers of trials for each of the 4 difficulty levels, pseudorandomly chosen across the 4 blocks. Stimuli were back-projected onto a screen located at the foot of the MRI bed using an LCD projector. Subjects viewed the screen via a mirror system located in the head coil and the start of each run was electronically synchronized with the MRI acquisition computer. Trial-by-trial feedback was not given, but the overall accuracy was presented at the end of each block. Behavioral responses were recorded with a 4-key fiber-optic response box (Resonance Technology, Inc.). Scanner noise was reduced with ear plugs, and head motion was minimized with foam pads. When necessary, vision was corrected using MRI-compatible lenses that matched the distance prescription used by the participant. The total scan time, including breaks and structural scans, was approximately 1 h 40 min.

### MRI scanning and data preprocessing

MRI was performed in a 3-T GE scanner at the at Duke Brain Imaging Analysis Center (BIAC). Structural MRI and DWI scans were followed by performing 4 fMRI runs of the DRAT task. The anatomical MRI was acquired using a 3D T1-weighted echo-planar sequence (matrix = 2562, TR = 12 ms, TE = 5 ms, FOV = 24 cm, slices = 68, slice thickness = 1.9 mm, sections = 248). In the fMRI runs, coplanar functional images were acquired using an inverse spiral sequence (64 × 64 matrix, time repetition [TR] = 2000 ms, time echo [TE] = 31 ms, field of view [FOV] = 240 mm, 37 slices, 3.8-mm slice thickness, 254 images). Finally, DWI data were collected using a single-shot echo-planar imaging sequence (TR = 1700 ms, slices = 50, thickness = 2.0 mm, FOV = 256 × 256 mm^2^, matrix size 128 × 128, voxel size = 2 mm^3^, b value = 1000 s/mm^2^, diffusion-sensitizing directions = 36, total images = 960, total scan time = 5 min).

Functional images were preprocessed using image processing tools, including FLIRT also from FSL, in a publicly available pipeline developed by the Duke Brain Imaging and Analysis Center (https://wiki.biac.duke.edu/biac:analysis:resting_pipeline). Images were corrected for slice acquisition timing, motion, and linear trend; motion correction was performed using FSL’s MCFLIRT, and 6 motion parameters estimated from the step were then regressed out of each functional voxel using standard linear regression. Images were then temporally smoothed with a high-pass filter using a 190s cutoff, and normalized to the Montreal Neurological Institute (MNI) stereotaxic space. White matter and CSF signals were also removed from the data, using WM/CSF masks generated by FAST and regressed from the functional data using the same method as the motion parameters. Spatial filtering with a Gaussian kernel of full-width half-maximum (FWHM) of 6mm was applied.

### Experimental Design and Statistical Analyses

#### Behavioral Analyses

Accuracy and response times (RTs) of correct DRAT trials were analyzed in terms of Set Size and Sorting Steps using linear mixed effects models, as implemented by R and lme4. **Set Size** had four levels, Very Easy, Easy, Medium, and Hard, which were defined based on data from the first session. Individual fitted accuracy functions, centered around each subject’s individual Criterion, and determination of their Starting Set Size (i.e. Set Size value corresponding to the Very Easy condition) are shown in **Figure 2A**. Across the sample of 41 participants, 12 had a Starting Set Size of 3; 19 had a Starting Set Size of 4; 9 had Starting Set Size of 5; and 1 had a Starting Set Size of 6. In all future references, Relative Set Size refers to the individually titrated load of four Set Sizes for each subject (beginning with their Starting Set Size, then +1 item, +2 items, and +3 items) quantified across four discrete levels (1-4), whereas Absolute Set Size refers to the original number of letters in an array.

**Figure 2.**
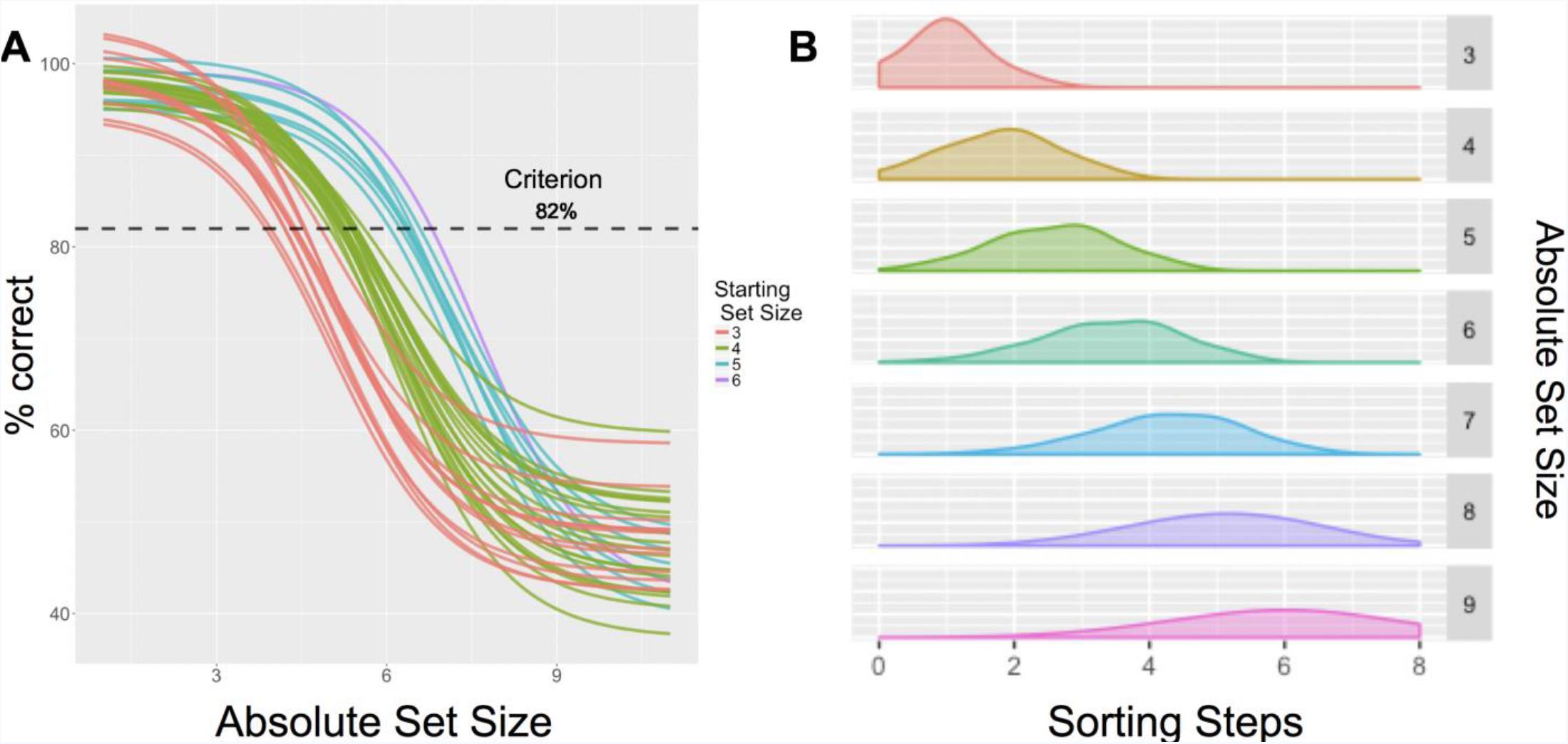
Interaction of Absolute Set Size and Sorting Steps. (**A**) Individually-titrated (Relative) Set Sizes were determined using sigmoid curves fitted to individual performance data, based on accuracy from screening. (**B**) Distribution of Sorting Steps across the Absolute Set Sizes in the current paradigm (i.e., before adjusting to 4-levels based on session 1 titration).

**Sorting Steps** is the minimum number of discrete changes required to transform the initial random letter array into the alphabetized array. The number of sorting steps was estimated using the minimum number of sorting operations calculated from four sorting algorithms (Golde et al., 2010): insertion, selection, merge insertion, and merge selection. *Insertion* consists of processing each letter one-by-one and inserting it into the correct alphabetized position. *Selection* consists of identifying the earliest letter in the alphabet and swapping it with the letter occupying the correct position. *Merge insertion* and *merge selection* are similar to insertion and selection, respectively, but they subdivide the letter array into two sub-arrays, sorting within each of them, and then combining the results. Assuming that participants used the most efficient strategy, sorting steps was calculated as the minimum number of reordering steps from among the four algorithms on each trial. Given the logical complexities in orthogonalizing Absolute Set Size and Sorting Step factors, letters were selected at random, approximating a normal distribution within each Absolute Set Size (**Figure 2B**).

Absolute Set Size and Sorting Steps were moderately correlated (r = 0.51). The distribution of Sorting Steps within each Absolute Set Size approximated a normal distribution within each level of Set Size (all Shapiro-Wilk tests, W = 0.81-.95), though increasing Set Size was naturally associated with a wider distribution in the number of Sorting Steps for that level (**Figure 2B**). To confirm that both Set Size and Sorting Steps had significant and independent effects on performance, linear (for RT) and logistic (for accuracy) regression analyses were conducted. In all subsequent analysis, Relative Set Size is used as the measure of Set Size to best standardize the level of difficulty across all subjects. RTs were analyzed only for correct trials using a linear restricted maximum likelihood model. Accuracy was analyzed using a binomial logistic model including all trials. For both models, Set Size and Sorting Steps were treated as fixed effects while individual subjects were treated as a random effect. In addition, for both RT and Accuracy models, the interaction term (Set Size by Sorting Steps) was tested in order to account for additional variance attributed to increasing Sorting Steps across the 4 levels of difficulty. In both models, R (R Core Team, 2012) and lme4 (Bates, Maechler & Bolker, 2012) were used to perform a linear mixed effects analysis; while Relative Set Size and Sorting Steps (with interaction term) were entered into the fixed effects model. Intercepts for subjects, as well as by-subject random slopes were entered for the random effects of Relative Set Size and Sorting Steps. Gender, age, and each subject’s Starting Set Size were also included to account for standardizing difficulty levels across subjects. Visual inspection of residual plots did not reveal any obvious deviations from homoscedasticity or normality. P-values were obtained by likelihood ratio tests of the full model with the effect in question against the model without the effect in question. There was no missing data, but participants failed to respond within the permitted 4-second time window on 1.6% of trials (79 out of 4920). These trials were excluded from all analyses.

#### fMRI analyses

A parametric approach was used to investigate how activity varied as a function of Relative Set Size and Sorting Steps. First-level voxel time-series analysis was carried out using general linear modeling (GLM) implemented in the FEAT toolbox of FSL. Fixed effects models were carried out to examine the parametric effects of Set Size and the number of sorting operations necessary to alphabetize each trial; separate events were modeled for the array presentation (duration: 3s), delay period (duration: 5s), and response (duration: subject response time), each with an onset at the beginning of the event. Weighted regressors during the delay period were used to model the difficulty associated with different WM operations. The first regressor increased linearly with the array’s Set Size to model the parametric increase in difficulty with increased letter load. The second weighted regressor reflected the minimum number of sorting steps needed on a given trial. Both of these parametric variables were orthogonalized to the non-parametric delay-period regressor, the trial period when maintenance and manipulation are likely to operate concurrently. Incorrect and non-response trials were modeled identically, but separately, and were not considered in the results below. Subsequent to individual-level models, random-effects analysis was performed on the parameter estimates of the parametric regressors (p < 0.005, cluster correction: z > 2.0).

### Cortical Parcellation

Before either structural or functional matrices were constructed, consistent parcellation scheme were established across all subjects and all modalities (DWI, fMRI) that reflect an accurate summary of the full connectome effects (Cocchi et al., 2014). Subjects’ T1-weighted images were segmented using SPM12 (www.fil.ion.ucl.ac.uk/spm/software/spm12/), yielding a grey matter (GM) and white matter (WM) mask in the T1 native space for each subject. The entire GM was then parcellated into 471 regions of interest (ROIs), each representing a network node by using a subparcellated version of the Harvard-Oxford Atlas, (Braun et al., 2015), defined originally in MNI space. The T1-weighted image was then nonlinearly normalized to the ICBM152 template in MNI space using fMRIB’s Non-linear Image Registration Tool (FNIRT, FSL, www.fmrib.ox.ac.uk/fsl/). The inverse transformations were applied to the HOA atlas in the MNI space, resulting in native-T1-space GM parcellations for each subject. Then, T1-weighted images were coregistered to native diffusion space using the subjects’ unweighted diffusion image as a target; this transformation matrix was then applied to the GM parcellations above, using FSL’s FLIRT linear registration tool, resulting in a native-diffusion-space parcellation for each subject.

### Structural connectivity

DWI data were analyzed utilizing FSL (https://fsl.fmrib.ox.ac.uk/fsl/fslwiki) and MRtrix (http://mrtrix.org) software packages. Data were de-noised with MRtrix, corrected with eddy current correction from FSL, and brain extraction was performed with both FSL and MRtrix, whereas bias-field correction was completed with MRtrix. Constrained spherical deconvolution (CSD) was utilized in calculating the fiber orientation distribution (FOD). This FOD was used along with the brain mask to generate whole brain tractography, with seeding done at random within the mask (Knuth, 1976; Beynel et al., in review). Relevant parameters regarding track generation are as follows: seed = at random within mask; step-size = 0.2 mm; 10,000,000 tracts. After tracts were generated, they were filtered using SIFT (spherical-deconvolution informed filtering of tractograms). This process utilizes an algorithm which determines whether a streamline should be removed or not based off of information obtained from the FOD, which improves the selectivity of structural connectomes by using a cost-function to eliminate false positive tracts (Yeh et al., 2016). Tracts were SIFTed until 1 million tracts remained. Prior to connectome generation, subject-specific MNI-space brains were created by an affine registration between the MNI T1 2mm brain template and b0s using FSL’s FLIRT. The MNI subject-specific brains then underwent another affine registration to the Harvard-Oxford 471 ROI template.

### Functional connectivity

Functional connection matrices representing task-related connection strengths were estimated using a correlational psychophysical interaction (cPPI) analysis used previously by us (Tzourio-Mazoyer et al., 2002) and others (Tournier et al., 2007) to estimate a whole-brain connectivity matrix that describes task-related interactions between brain regions. Briefly, the model relies on the calculation of a PPI regressor for each region (or node), based on the product of that region’s timecourse and a task regressor of interest, in order to generate a term reflecting the psychophysical interaction between the seed region’s activity and the specified experimental manipulation.

#### Network definition

In the current study, subjects’ T1-weighted images were segmented using SPM12 (www.fil.ion.ucl.ac.uk/spm/software/spm12/), yielding a grey matter (GM) and white matter (WM) mask in the T1 native space for each subject. The entire GM was then parcellated into 471 regions of interest (ROIs), each representing a network node by using a subparcellated version of the Harvard-Oxford Atlas, (Tournier et al., 2004), defined originally in MNI space. The T1-weighted image was then nonlinearly normalized to the ICBM152 template in MNI space using fMRIB’s Non-linear Image Registration Tool (FNIRT, FSL, www.fmrib.ox.ac.uk/fsl/). The inverse transformations were applied to the HOA atlas in the MNI space, resulting in native-T1-space GM parcellations for each subject. Next, the convolved task regressors from the univariate model described above were used as the psychological regressor, which were originally coded as either a) the unmodulated (weight = 1) delay for each trial, b) the Set-Size-modulated delay regressor (range = 1-4), or c) the Sorting Operations-modulated delay regressor (range = 0-7); all regressors are mean-adjusted in FSL. Additional psychological regressors were modeled on the onsets for encoding (i.e., letter array) and response (i.e., cue) periods, but were not used in the connectivity analysis. The delay-period regressors were each multiplied with two network timecourses for region *i* and *j*. Partial correlations 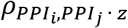 were then computed by removing the variance *z*, which includes both the psychological regressor and the time courses for regions *i* and *j*, as well as constituent noise regressors including 6 motion parameters and noise regressors coding for the concurrent signal in white matter and CSF during each run. In order to compare equally reliable estimates of connectivity delineated by either Set Size or Sorting Steps, the distribution of Sorting Steps within each individual from 0-7 to 1-4 level was interpolated, such that an equal number of trials were used to estimate connectivity values in each parameter. This cPPI analysis resulted in 8 separate output matrices, comprising connectivity delineated by Set Size (4 levels), or Sorting Steps (also 4 levels). Task-related connectivity was estimated from the resulting output matrices; negative connections were included in these analyses, as they may inform important, explicit interpretations about how networks may be segregated (Yeh et al., 2016). Graph metrics, including modularity (describing the modular organization of the whole-brain graph) and strength (describing a sum of the connectivity strengths for each node) were computed using the Brain Connectivity Toolbox as described previously (Davis et al., 2017) and, when appropriate, summed across all nodes within a task-related network.

Maintenance and Manipulation networks were defined by using both functional and structural information. First, parametric univariate activity from voxelwise maps was averaged within individual regions of interest (ROI) within the 471-ROI Harvard-Oxford brain atlas, and ranked by mean z-score. This information was used to identify the top 5% nodes for each parametric effect. Both networks were constructed with equal numbers of nodes, in order to ensure that the main network metrics (within-or between-network correlations, see below) were not biased by the number of regions contributing to that aggregate measure. Each ROI was ranked by its mean parametric effect z-score and the top 5% of nodes were classified as either Maintenance or Manipulation network nodes. Lastly, structural connectivity information (FA of each pairwise connection) between all network nodes (5% of 471 = 23 Maintenance nodes, 23 Manipulation nodes) was assessed for both within- and between-network connection strength.

#### Network segregation and reorganization measures

Lastly, in order to summarize the more system-wide behavior of the two task-related networks, two derived measures of overall network organization were calculated. First, a previously reported (Fornito et al., 2012) measure of system segregation was used. This measure was calculated as the difference between the mean magnitudes of between-system correlations from the within-system correlations as a proportion of mean within-system correlation.

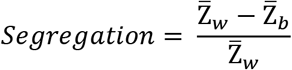

Where Z̄_*w*_ is the mean r-values between nodes of one partition, module, or system (similar to within-module degree or WMD), and Z̄_*b*_ is the mean of r-values between nodes of separate partitions (similar to between-module degree or BMD, Tzourio-Mazoyer et al., 2002). Accordingly, values greater than 0 reflect relatively lower between-system correlations in relation to within-system correlations (i.e., stronger segregation of systems), and values less than 0 reflect higher between-system correlations relative to within-system correlations (i.e., diminished segregation of systems).

Second, a network reconfiguration measure was developed to describe the similarity in functional connectivity across the task conditions. While the segregation measure above is descriptive of network behavior at discrete levels of difficulty, network reconfiguration describes the overall similarity between task conditions, i.e., between network states. Network reconfiguration represents a direct comparison between network states, and in this case represents an average of the correlation values between all functional connection matrices for a given subject.

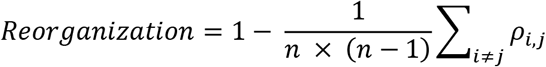

Where *n* is the number of states (e.g., 4 in this case), and *ρ*_*i*,*j*_ represents the Spearman’s correlation between the complex functional connectivity profiles representing two brain states *i* and *j* (e.g., functional connectivity matrices representing Easy and Medium difficulty levels in this case). Thus, highly correlated matrices represent low reconfiguration (closer to 0), while weakly correlated matrices represent high reconfiguration across task conditions (closer to 1). Given the explicit hypotheses concerning segregation and integration of the putative Maintenance and Manipulation networks, reconfiguration within a subset of connections that describe a) connections within the Maintenance network, b) connections within the Manipulation network, and c) connections between both networks were examined.

## Results

### Behavioral results

Figure 3 presents accuracy and RT data. Based on likelihood ratio tests of the full model with a null model removing the relevant term (**Table 1**), both Relative Set Size (4 load levels titrated to individual performance) and Sorting Steps made significant and distinct contributions to both accuracy and RTs. Specifically, the binary logistic regression of accuracy revealed a significant effect of Relative Set Size (X^2^ = 80.07, p = 2.2e-14) and Sorting Steps (χ2 = 22.14, p = 2.5e-6), as well as a significant Relative Set Size by Sorting Steps interaction (X^2^ = 12.35, p = 4e-4). The linear mixed effects regression applied to RT data revealed a similar pattern of findings, such that both Relative Set Size (χ^2^ = 45.73, p = 1.4e-11), Sorting Steps (X^2^ = 12.39, p = 4.3e-4), and their interaction (X^2^ = 10.66, p = 1.1e-3) demonstrated significant effects. Effects of Gender and Starting Set Size were nonsignificant in both models (p>0.05), which is not surprising given the inclusion of intercepts for subjects, as well as by-subject random slopes for the effect of Relative Set Size and Sorting Steps. These findings therefore support the approach of using these two measures to disentangle maintenance and manipulation WM mechanisms.

**Figure 3.**
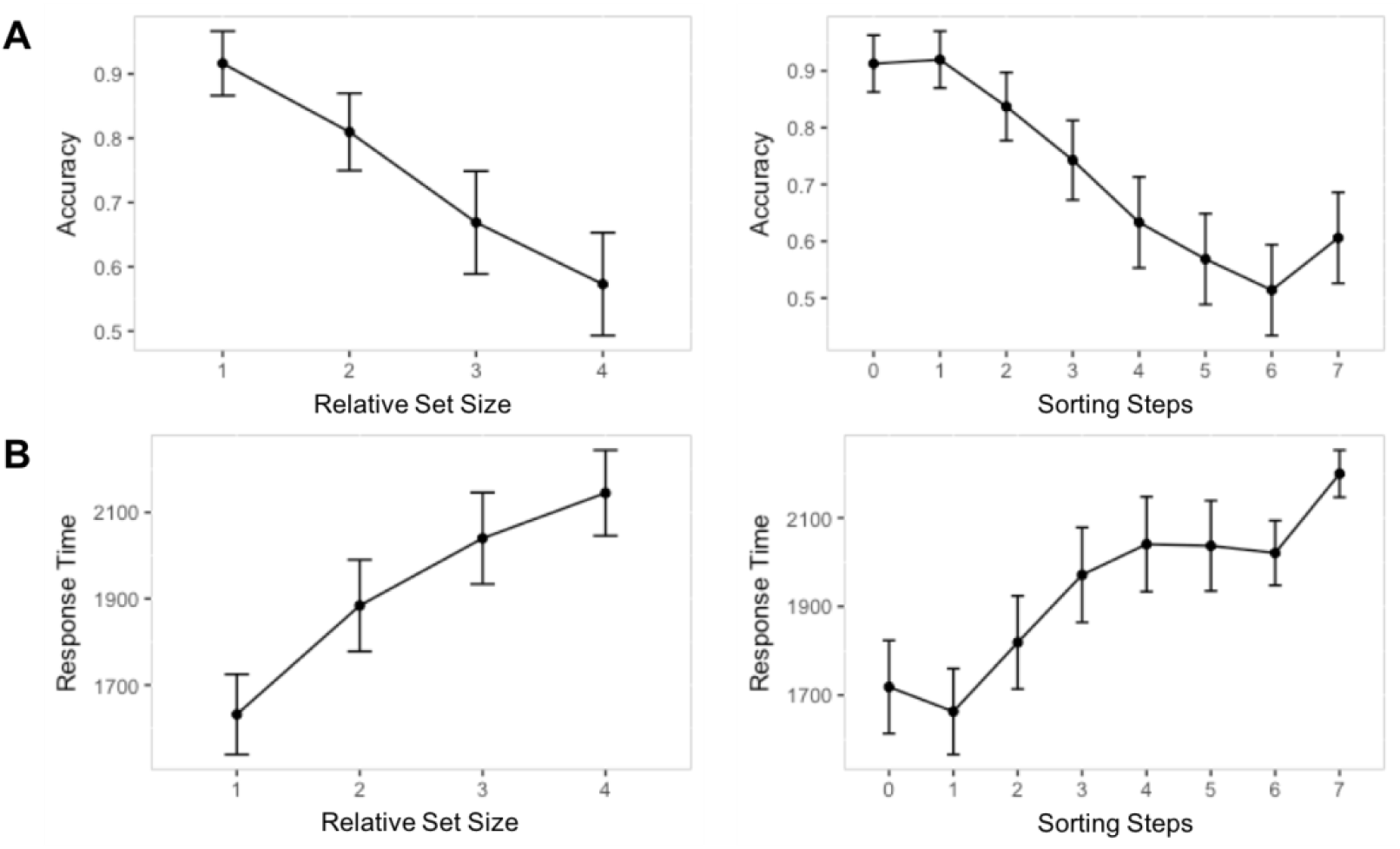
Mean values and standard error across subjects for accuracy (**A**) and RTs (**B**) across Relative Set Size, reflecting the number of items to be retained in WM across a 5s delay (adjusted across subjects to 4 levels), and Sorting Steps, reflecting the number of sorting operations required to alphabetize a given letter array. Note: Statistical significance was determined by linear mixed-effects models.

**Table 1.** ANOVA of factors affecting accuracy and RTs

| Effect | Estimate | Std.Error | $\chi^2$ Value | Pr > \|t\| |
| --- | --- | --- | --- | --- |
| Accuracy |  |  |  |  |
| Intercept | 4.10 | 0.32 |  |  |
| Set Size | -0.94 | 0.11 | 77.43 | 2.2e-14 |
| Sorting Steps | -0.49 | 0.10 | 23.46 | 2.1e-6 |
| Set Size * Sorting Steps | 0.11 | 0.03 | 12.35 | 0.0004 |
| Reaction times |  |  |  |  |
| Intercept | 1129.17 | 206.66 |  |  |
| Set Size | 284.36 | 37.05 | 45.73 | 1.4e-11 |
| Sorting Steps | 89.22 | 25.22 | 12.39 | 0.0004 |
| Set Size * Sorting Steps | -29.14 | 8.86 | 10.66 | 0.0031 |

Note: $\chi^2$ statistics and p-values were obtained by likelihood ratio tests of the full model with the effect in question against the model without the effect in question.

### fMRI results

#### Univariate activity

Univariate analyses were used to identify regions where delay-period activity increased parametrically as a function of Relative Set Size or Sorting Steps. As shown by **Figure 4A** and **Table 2**, models with *concurrent* parametric regressors show that Relative Set Size was associated with increased activity in bilateral PFC (including the middle and inferior frontal gyri—MFG and IFG), ventral parietal cortex (VPC), and the anterior cingulate cortex (ACC), whereas Sorting Steps were associated with activations in superior parietal lobule (SPL), ACC, the posterior cingulate cortex (PCC), the superior temporal gyrus (STG), and the hippocampus. Comparison of non-competing parametric maps at the single subject level confirmed that both maintenance and manipulation parameters elicited activity in overlapping middle-cingulate regions.

**Figure 4.**
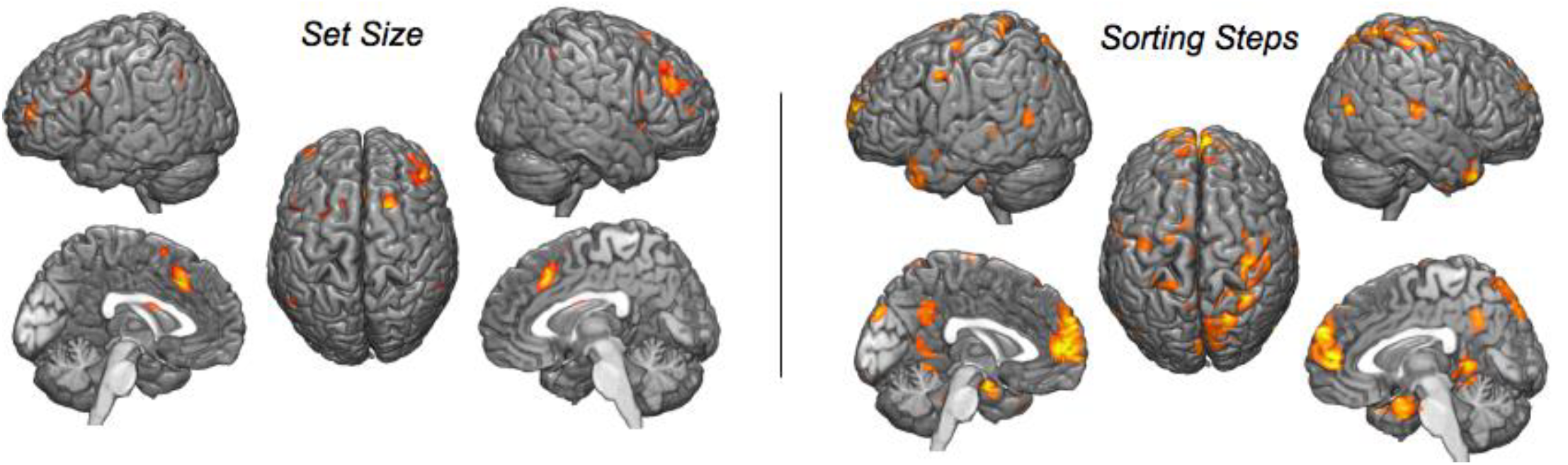
Parametric effects during the Delay period of Set Size, reflecting the number of letters in the encoding array, and Sorting Steps, reflecting the minimum number of reorganizing operations required to alphabetize an array. Images thresholded at p < 0.005, cluster correction FWE p < 0.05

**Table 2.**
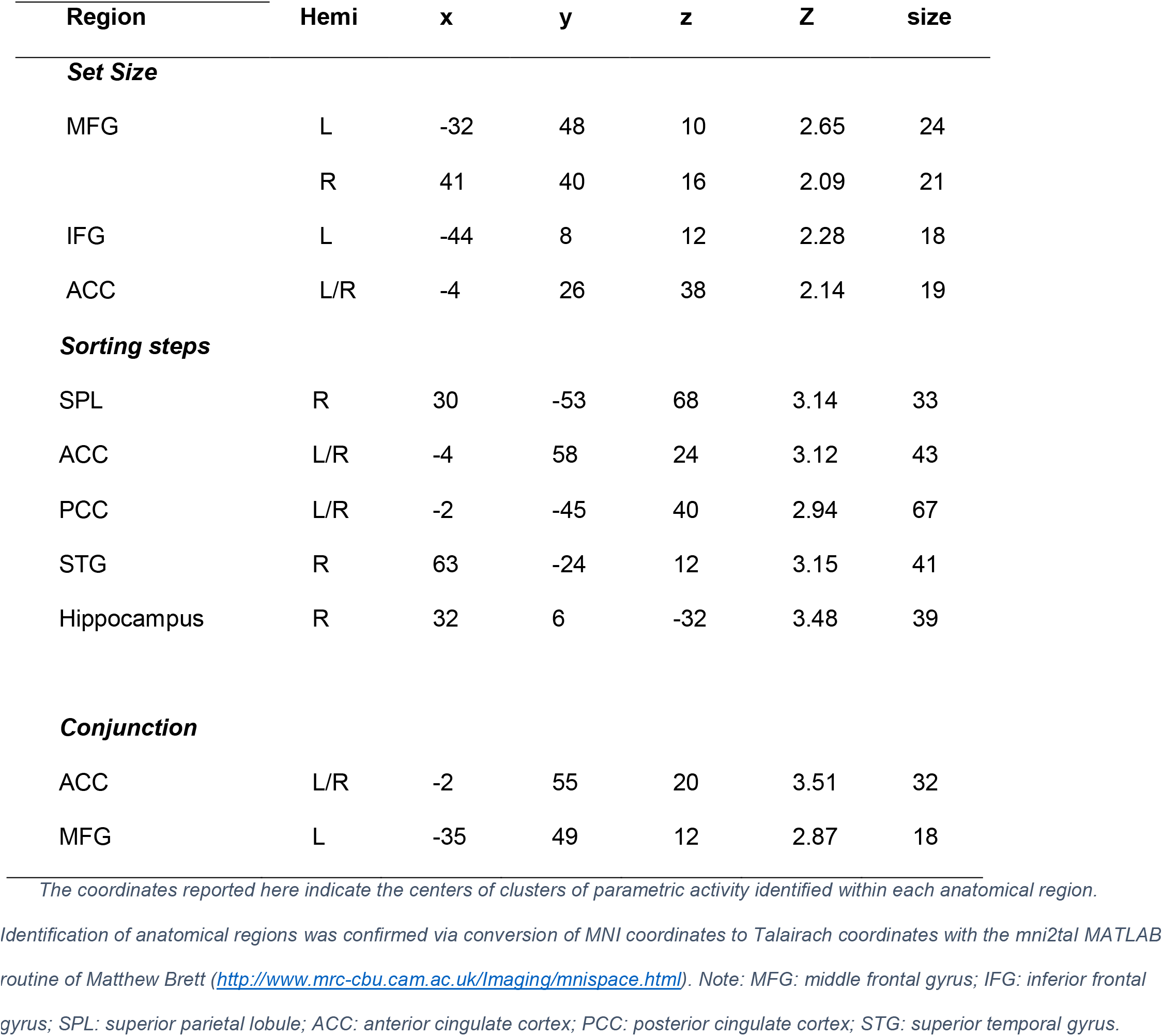
Parametric fMRI effects of Set Size and Sorting Steps

| Region | Hemi | x | y | z | Z | size |
| --- | --- | --- | --- | --- | --- | --- |
| **Set Size** |  |  |  |  |  |  |
| MFG | L | -32 | 48 | 10 | 2.65 | 24 |
|  | R | 41 | 40 | 16 | 2.09 | 21 |
| IFG | L | -44 | 8 | 12 | 2.28 | 18 |
| ACC | L/R | -4 | 26 | 38 | 2.14 | 19 |
| Sorting steps |  |  |  |  |  |  |
| SPL | R | 30 | -53 | 68 | 3.14 | 33 |
| ACC | L/R | -4 | 58 | 24 | 3.12 | 43 |
| PCC | L/R | -2 | -45 | 40 | 2.94 | 67 |
| STG | R | 63 | -24 | 12 | 3.15 | 41 |
| Hippocampus | R | 32 | 6 | -32 | 3.48 | 39 |
| Conjunction |  |  |  |  |  |  |
| ACC | L/R | -2 | 55 | 20 | 3.51 | 32 |
| MFG | L | -35 | 49 | 12 | 2.87 | 18 |

The coordinates reported here indicate the centers of clusters of parametric activity identified within each anatomical region. Identification of anatomical regions was confirmed via conversion of MNI coordinates to Talairach coordinates with the mni2tal MATLAB routine of Matthew Brett (http://www.mrc-cbu.cam.ac.uk/Imaging/mnispace.html). Note: MFG: middle frontal gyrus; IFG: inferior frontal gyrus; SPL: superior parietal lobule; ACC: anterior cingulate cortex; PCC: posterior cingulate cortex; STG: superior temporal gyrus.

The strength of these unique effects is surprising, given the moderate collinearity between Set Size and Sorting Steps noted above. To investigate possible overlaps between the parametric effects of Set Size and Sorting Steps, whole-brain conjunction analysis was performed at the subject level, using parametric fMRI models with *either* Set Size or Sorting Steps (but not both regressors). Significant overlapping voxels were observed only in mid-cingulate cortex and anterior SFG, indicating that these regions are sensitive to both maintenance and manipulation.

Furthermore, the model fit for each ROI was examined to infer explicit evidence for collinearity between the convolved parametric Set Size and Sorting Step regressors. In order to test explicitly for the nature of the collinearity between these terms, the average Variance Inflation Factor (VIF) was calculated across runs, for each ROI. A VIF is the ratio of variance in a model with multiple terms, divided by the variance of a model with one term alone (Braun et al., 2012); large VIFs are a measure of multicollinearity, and thus a test of the specific parametric factors can help validate whether these terms carry unique information. VIFs were calculated on the full first-level models (i.e., each run), comprising convolved regressors for all parametric and nonparametric events; VIFs for parametric Set Size and Sorting Steps effects were then averaged across runs. These analyses revealed that the VIF for both Set Size (2.58, SD = +/- 0.48 across subjects) and Sorting Steps (VIF = 2.49, SD = +/- 0.48 across subjects) remained well within established guidelines for the VIF (general VIF < 5; see Rubinov and Sporns, 2010; Chan et al., 2014).

#### Network analyses

The network-level analyses are organized into 3 stages: network identification and validation, basic network description, and segregation & reconfiguration analysis. These analyses began by identifying Maintenance and Manipulation networks, by relying on both functional and structural information to define and validate the task-based connectivity approach. These networks were constructed with equal numbers of nodes, in order to ensure that the main network metrics (within-or between-network correlations, see below) were not biased by the number of regions contributing to that aggregate measure. First, masked parametric univariate activity (**Figure 5A**) with the 471-ROI Harvard-Oxford brain atlas was used in order to identify the top 5% nodes (n = 23) for each parametric effect (**Figure 5B**), as determined by the z-statistic from the parametric map within a given ROI/node; no overlapping nodes were found. To ensure an equal number of ROIs in the two networks, each ROI was ranked by its mean z-score in parametric analyses and identified the top 5% of nodes (a more liberal top-10% or top-20% threshold [n = 46, 92 nodes in each network] also revealed no overlap in networks). The Maintenance (blue) and Manipulation (blue) networks are visualized both as the nodes and as the connections between these nodes in **Figure 6A**.

**Figure 5.**
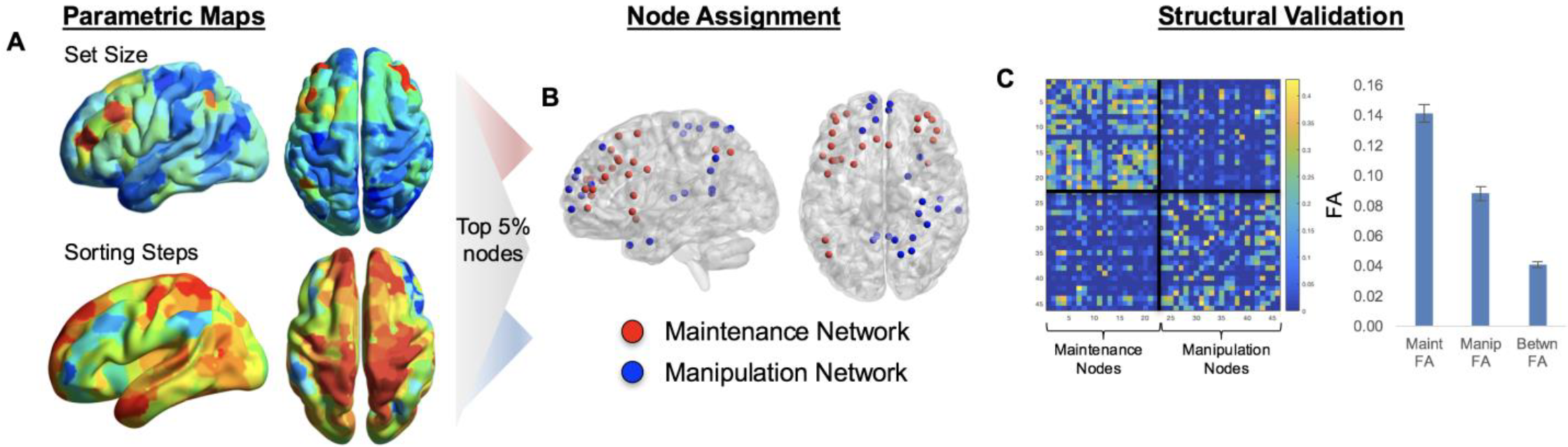
Converting univariate information into multivariate topology. Thresholded parametric maps (Fig. 4), using average responses within all voxels in each ROI in the HOA471, were used in order to identify regions responsive to Set Size or Sorting Steps (**A**). (**B**) The top 5% nodes of each network were then assigned to either Maintenance or Manipulation networks, based on the parametric effect (z-score) within these nodes (p < 0.005).(**C**) Structural network connectivity is stronger within than between networks, helping to validate the task-based network parcellation. Independent of any functional information, nodes selected within the Maintenance or Manipulation networks showever greater connectivity than between the two putative task-related networks.

**Figure 6.**
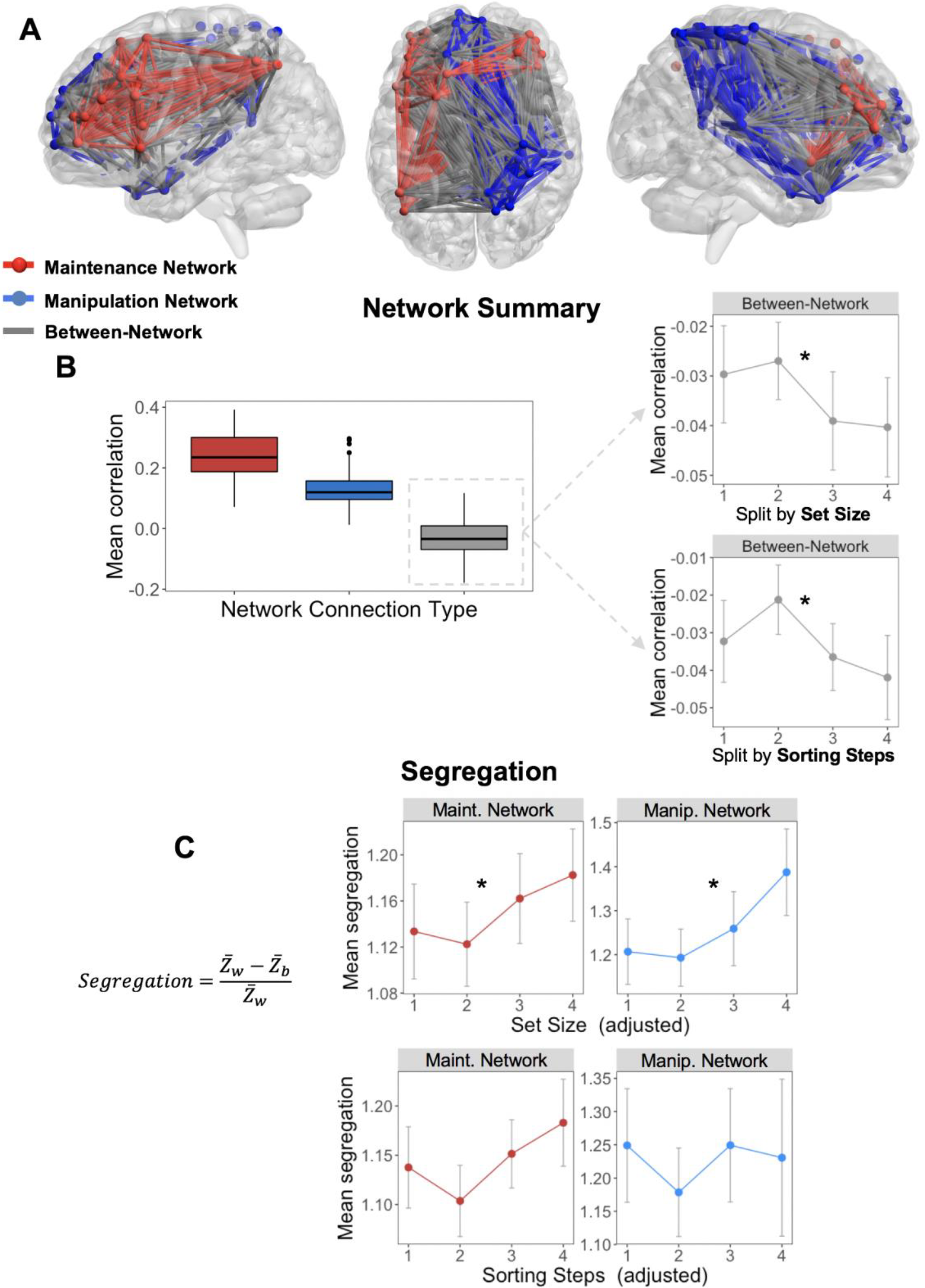
Parametric effects of Set Size and Sorting Steps on nodes within the Maintenance and Manipulation networks, respectively. (**A**) Schematic describes the organization of either within (red, blue) or between (grey) network connections. The Maintenance network is generally described as a bilateral frontal network (with specific connections to left IPS), while the Manipulation network connects larely right SPL and midline frontal regions. (**B**) Within the Maintenance and Manipulation networks, within-network connections remain consistently positive, while all between-network connections are negative, with a negative trend with increasing Set Size or Sorting Steps. (**C**) Both Maintentance and Manipulation networks became increasingly segregated with increasing Set Size, suggesting that the negative relationship between these two networks was behaviorally meaningful. Note: Statistical significance was determined by linear mixed-effects models, which may not be reflected in the averages and standard errors displayed here.

##### Structural network validation

Before analyzing functional within- and between-network connectivity, averaged across the putative task-related networks, patterns of structural connectivity between nodes was examined in order to test the validity of the task-based node definitions. If these networks form reliable task-based parcellations, structural network connectivity should be weaker between-networks than within-networks. Consistent with this idea, structural connection strength (measured using fractional anisotropy) was weaker between-networks than within-network, in either Maintenance (*t_28_* = 20.5; p = 2.2e-18) and Manipulation (*t_28_* = 12.7; p = 3.5e-13) networks (**Figure 5C**). This result suggests a structural basis for functional connectivity patterns within each task-related network, and points to a clear structural hurdle to between-network connections. While these effects may be at least partially due to greater mean distance between nodes (Maintenance network: 57.1mm; Manipulation network: 73.7mm; between-network: 82.4mm), this difference is not incompatible with community membership (regions closer together are often more likely to form coherent neurocognitive networks). Thus, subsequent network analysis results are characterized in terms of two discrete networks, the "Maintenance network" and "Manipulation network". While we have demonstrated that this task-based community assignment has both functional and structural foundations, we do not assume that the same Maintenance and Manipulation networks operate for every particular WM paradigm and stimulus type.

##### Effects of Set Size and Sorting Steps on summary measures of functional network connectivity

Next we return to our two principle measures of Maintenance and Manipulation functions, and examine the effects of increasing Set Size or Sorting Steps, respectively, on within-and between-network connections (**Figure 6A**) were examined in the same discrete Maintenance and Manipulation networks defined above. Here, two reliable patterns were found that helped explain how increasing computational complexity in the behavioral domain manifests as a more segregated cortical system in which local networks predominate over more global connectivity. As illustrated by **Figure 6B** a significant main effect of Network Connection Type on connectivity (i.e., mean correlation value, F_1,39_ = 215.23, p < 0.001) was found, such that the mean correlation values were stronger in the Maintenance and Manipulation networks than between the networks. When difficulty was split by Set Size, within-network connectivity in both the Maintenance and Manipulation networks was consistently positive (one-sample *t* collapsing across levels were 5.31 and 4.43, respectively, both p < 0.001), as may be expected for networks defined by their task-relatedness. Splitting these same networks by Sorting Steps elicited similar effects. Chi-squared tests accounting for subject-level differences in mean connectivity demonstrated no effect of difficulty on within-network connectivity in either Maintenance or Manipulation network, whether difficulty was defined by Set Size (χ2 = 0.3, χ2 = 0.5, respectively for each network, both p > 0.1) or by increasing number of Sorting Steps (χ^2^ = 0.6, X^2^ = 0.2, respectively, both p > 0.1). This result suggests that the connectivity between nodes within each network was consistent across all levels of difficulty, and that any difficulty-related changes are driven largely by between-network connections.

Interestingly, in contrast with the positive within-network connections (range for Maintenance network: r = 0.22-0.25, Manipulation network: r =0.11-0.13), between-network correlations were consistently negative (mean r across levels = −0.04; one-sample t-test collapsing across levels: t_40_ = −3.57, p = 4.62e-3). Furthermore, the mean connectivity between networks demonstrated a negative decline with increasing Set Size (χ^2^ = 3.81, p = 4.5e-2) or increasing number of Sorting Steps (X^2^ = 3.51, p = 5.4e-2), indicating that the correlation between nodes in these two task networks declines linearly with increasing complexity, signifying a behaviorally meaningful relationship.

##### Network re-organization and its behavioral consequence

To examine system-level organization, two derived measures of overall network organization were calculated. Segregation, which describes the difference between within- and between-network correlations as a proportion of mean within-system correlation, is defined by the segregation coefficient, a node-level measure describing the degree to which local nodes become more connected to other local nodes within a network compared to nodes outside the local network (**Figure 6C**). Within both the Maintenance and Manipulation networks, the segregation coefficient showed a clear linear increase with increasing Set Size (χ^2^ = 4.53, X^2^ = 4.48, respectively, both p < 0.05), further supporting the idea that the global organization tended towards increasingly segregated network nodes. In contrast, increasing Sorting Steps did not elicit the same effect in either the Maintenance or Manipulation networks (χ^2^ = 2.34, X^2^ = 0.1, respectively, both p > 0.1), suggesting that the segregation effect was driven by changes in Set Size.

Lastly, network reconfiguration was analyzed using a summary statistic that describes the individual differences in network reconfiguration across the task conditions (**Figure 7A**). Here, it was found that network reconfiguration was greater in connections between Maintenance and Manipulation networks than within either task network alone (t = 9.84, p = 4.1e-12; t = 11.10, p = 1.2e-13, respectively; see marginal rug plots in **Figure 7B**). Furthermore, network reconfiguration in these between-network connections was predictive of subjects’ individual Criterion for the WM task, which describes the idealized 82% level of behavioral performance (r_39_ = 0.39, p = 0.012), while within-network reconfiguration was not (r_39_ = 0.17 and 0.30 for Maintenance and Manipulation networks, respectively, both p > 0.05). Results were similar when splitting networks by the number of Sorting Steps, with a slight increase in the correlation between between-network reconfiguration and Criterion (r_39_ = 0.41, p = 0.007). The direction of these effects demonstrates that individuals with higher working memory capacity have greater changes in between-network functional connectivity in response to increasingly difficult task conditions, suggesting that network reconfiguration in working-memory related regions is adaptive to task demands.

**Figure 7.**
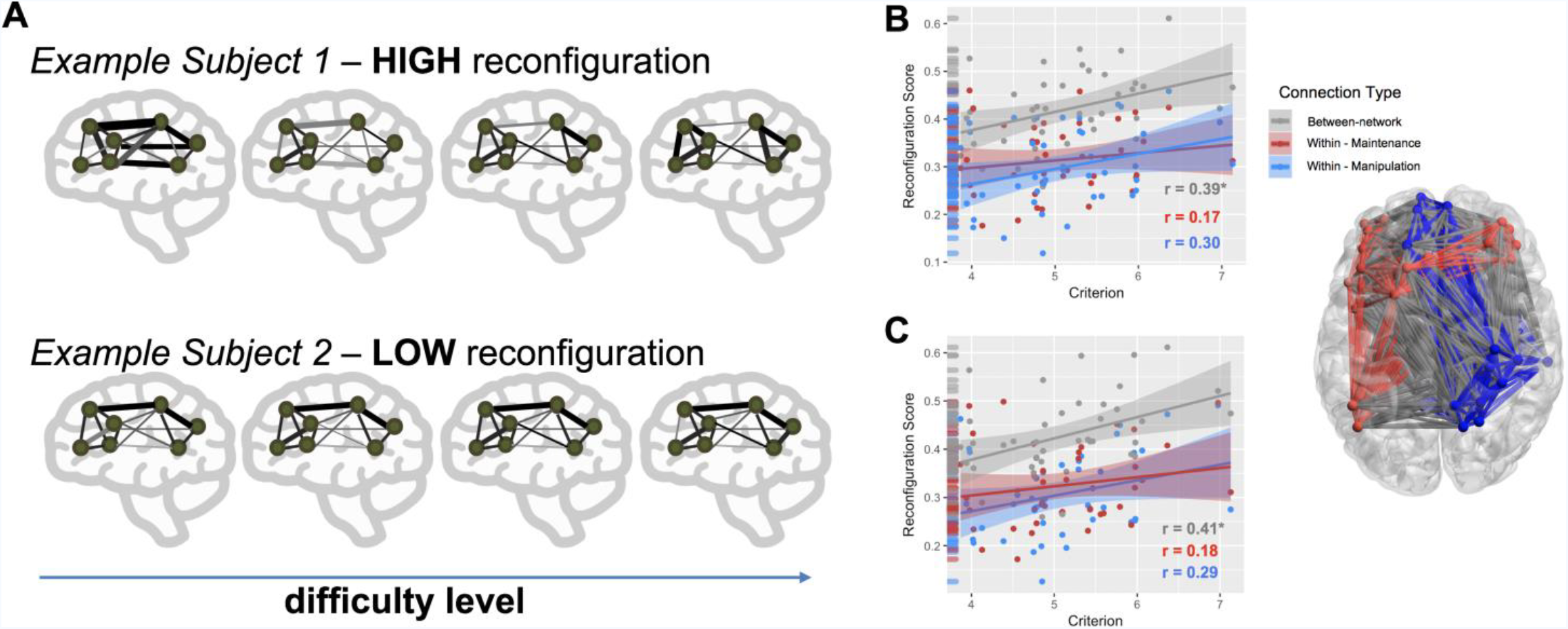
Network Reconfiguration. (**A**) While the segregation measure above is descriptive of network behavior at discrete levels of difficulty, network reconfiguration describes the overall similarity between task conditions, i.e., between network states. Network reconfiguration represents a direct comparison between network states, and in this case represents an average of the correlation values between all functional connection matrices for a given subject. (**B**) Network reconfiguration was higher in between-than within-network connections (B), and predictive of individual differences in working memory ability (i.e., Criterion).

## Discussion

Going beyond previous fMRI studies on WM maintenance versus manipulation, the current study investigated these processes using a novel behavioral paradigm in which maintenance and manipulation are assessed by indexing maintenance in terms of Set Size (number of letters) and manipulation in terms of Sorting Steps (number of sorting operations to alphabetize a letter array). The study yielded three main findings. First, it was found that Set Size and Sorting Steps made significant and independent contributions to accuracy and RTs, supporting the distinction between maintenance and manipulation. Second, maintenance and manipulation recruited distinct frontal-parietal patterns of univariate activity: maintenance was associated with a bilateral fronto-parietal network, as typical in WM tasks, whereas manipulation was associated with greater activity in the right SPL, a region associated with symbolic computations. Third, summary measures of the functional connectivity between the Maintenance and Manipulation networks demonstrated a negative association which increased with task demand, suggesting the action of a protective mechanism against interference of the cognitive operations within the two networks. These three main findings are discussed below.

### WM maintenance and manipulation are dissociable in behavior

The first goal of this study was to provide evidence that distinct, concurrent processes underlie basic working memory operations. To the authors’ knowledge, this is the first study in which WM maintenance and manipulation have been investigated during the same task. Moreover, it is also the first instance in which WM manipulation has been linked to a specific measure of the computation required by the task, namely the alphabetization of letters which requires individuals to sort letters into an ordered array. Here, the number of sorting steps was quantified using established algorithms from the computer science literature (insertion sort, selection sort, etc.). Although Set Size and Sorting Steps measures were correlated, it was possible to disentangle their effects on WM behavior (accuracy and RTs) and brain activity. In particular, regression analyses showed that both these measures significantly involved in WM performance, but their effects were independent, consistent with the idea that neural mechanisms of maintenance and manipulation are dissociable.

### Distinct univariate brain activity for concurrent maintenance and manipulation operations

Satisfying the second goal of this study, strong evidence was found for concurrent univariate brain activity tracking separate maintenance and manipulation operations during the WM task. Set Size was associated with activations in bilateral frontal and parietal regions, whereas Sorting Steps was associated with selective recruitment of a right SPL region, as well as activations in ACC, STG, and hippocampus. Below, we consider the two sets of regions associated with maintenance and manipulation.

The finding that maintenance was associated with bilateral fronto-parietal is consistent with fMRI evidence linking these regions to WM capacity (Kraha et al., 2012).. Within PFC, the current results linked maintenance to dorsolateral PFC (DLPFC). In previous studies, maintenance—as indexed by Set Size or load—has been sometimes linked to ventrolateral PFC activity (Braver et al., 1997; Mumford et al., 2015), but this linked has been challenged (De Pisapia and Braver, 2008). Also, several studies have linked DLPFC to manipulation, not to maintenance (Ranganath et al., 2004b; Ranganath et al., 2004a; Postle et al., 2006; Libby et al., 2014). However, these studies investigated maintenance and manipulation in separate tasks. A problem with this general isolated approach to maintenance and manipulation is that, compared to maintenance tasks (e.g., holding letters in order), manipulation tasks (e.g., alphabetizing) involve not only greater manipulation (e.g., sorting) but also greater maintenance (e.g., holding both original and reorganized letter sequences), as well as interference that may arise between the two processes. Thus, the differential involvement of DLPFC in manipulation tasks could reflect increased maintenance demands, rather than specific manipulation operations.

The finding that Sorting Steps during alphabetization was associated with right SPL is intriguing because this is not a region typically associated with WM manipulation. However, the link between Sorting Steps and right SPL is consistent with the role of this region in symbol computation. For example, activations in right SPL have been reported in almost every neuroimaging study of numerosity, including tasks primarily involved in basic quantity processing (Postle et al., 2006; Schedlbauer et al., 2014), as well as more precise number processing and numerical operations (Overath et al., 2015; Hullett et al., 2016). However, the role of SPL is not limited to number-based operations. There is also evidence of this region being similarly activated across tasks manipulating both numbers and letters, which may be the result of one or more underlying computational processes shared across domains of symbol manipulation (Cantlon et al., 2006). Thus, although right SPL is more commonly associated with number processing, its engagement in this task is likely the result of a more general process involved in all symbol-based computation.

In addition to right SPL, the number of Sorting Steps were also associated with STG and hippocampus. The hippocampus is commonly associated with successful spatial WM (Wylie et al., 2004), and STG is often related to auditory processing (Christodoulou et al., 2001; Eldreth et al., 2006). In this task, the activation in hippocampus may be associated with the mental rearrangement of the letters in space, and the STG with imagery of the letters’ while alphabetization was taking place. The involvement of this constellation of regions therefore suggests that symbol computation and rehearsal may be an intrinsic part of working memory manipulation.

### WM Maintenance and Manipulation networks are negatively associated

The third goal of this study was to investigate whether maintenance and manipulation differ not just on univariate activity but also on network interactions, as measured using graph theory. It was found that nodes in the Maintenance and Manipulation networks were consistently segregated across task conditions, such that summary measures of between-network connectivity were consistently negative. In addition to this general negative correlation, nodes between the two networks showed a consistent linear decrease in connectivity with increasing number of items, and increasing segregation with increasing task difficulty. These results suggest that these two dissociable networks maintain segregated, but significant interactions in order to dissociate the cognitive processes. The increasingly negative relationship with increasing difficulty suggests that these networks become more segregated to combat the interference of these processes as cognitive demand increases.

Recent empirical work has begun to focus on how selective network properties change between increasingly complex task conditions (Piazza et al., 2007; Stevens et al., 2012; Park et al., 2014), and how such changes in the modular structure of functional brain networks relate to behavior. While changes in modular structure in response to task difficulty have been observed now in a number of studies, one discrepancy is in the direction of the effect: both increases (Gruber et al., 2001) and decreases (Braun et al., 2015; Hearne et al., 2017) in modularity have been reported with increasing task complexity. The discrepancy in these findings may be related to the use of global network variables (e.g., global efficiency) and global network assignments (e.g., default mode network, salience network, etc.), both of which may conflate task-specific operations with operations or regions unrelated to the task at hand. In this context, the task-specific network approach used here first identified specific cortical nodes with relevance to the task, and then offered a clear mechanistic demonstration that the interaction between these systems is modulated by the task demands. This is supported by the increasing network segregation with task difficulty, suggesting that the maintenance of the letter arrays in working memory is increasingly protected from the interference generated by the manipulation of this information. Nonetheless, one result which unites these findings is that the degree to which individual subjects are able to make flexible adjustments in functional network structure is a strong predictor of behavioral performance (Cole et al., 2013; Simony et al., 2016). In particular, individuals who showed greater dynamic reconfiguration across maintenance or manipulation levels had better working memory capacity (as estimated by the subject-level Criterion values). Furthermore, this effect was limited to the reconfiguration of between-network connections (**Figure 7**), highlighting the key role of internetwork connectivity in mediating flexible behaviors. How such modular architecture supports the dynamic integration of many high-level cognitive functions remains far from understood, but the present results highlight the importance of task-related connectivity in WM maintenance and manipulation.

#### Conclusions

The current study presents evidence and arguments for two distinct cognitive functions supporting WM processing during short delays. We examined evidence for significant and independent contributions of Set Size and Sorting Steps in a WM alphabetization task, contributions reflecting Maintenance and Manipulation operations, respectively. These dissociable operations were mirrored in the univariate fMRI results, such that distinct patterns of bilateral fronto-parietal (Maintenance) and right-lateralized SPL (Manipulation) networks were activated. Lastly, we found that connectivity between these networks was increasingly segregated as difficulty increased, and that this effect was positively related to individual WM ability. This analysis therefore suggests the action of a protective mechanism against interference of the cognitive operations within dissociable components of the WM system.

## Acknowledgments

The authors would like to thank Al Erkanli for assistance with statistical analyses.

## Notes

**Conflicts of Interest:** The authors declare no competing financial interests.

## References

Baddeley A (2000) The episodic buffer: a new component of working memory? Trends in cognitive sciences 4:417–423.

Beynel L, Davis SW, Crowell CC, Hilbig SA, Lim W, Nyguyen D, Peterchev AV, Luber B, Lisanby SH, Cabeza R, Appelbaum LG (in review) Effects of 5Hz rTMS to dorsal lateral prefrontal cortex on working memory manipulation abilities, and potential predictors of rTMS effect.

Braun U, Schafer A, Walter H, Erk S, Romanczuk-Seiferth N, Haddad L, Schweiger JI, Grimm O, Heinz A, Tost H, Meyer-Lindenberg A, Bassett DS (2015) Dynamic reconfiguration of frontal brain networks during executive cognition in humans. Proceedings of the National Academy of Sciences of the United States of America 112:11678–11683.

Braun U, Plichta MM, Esslinger C, Sauer C, Haddad L, Grimm O, Mier D, Mohnke S, Heinz A, Erk S, Walter H, Seiferth N, Kirsch P, Meyer-Lindenberg A (2012) Test-retest reliability of resting-state connectivity network characteristics using fMRI and graph theoretical measures. NeuroImage 59:1404–1412.

Braver TS, Cohen JD, Nystrom LE, Jonides J, Smith EE, Noll DC (1997) A parametric study of prefrontal cortex involvement in human working memory. NeuroImage 5:49–62.

Bunge SA, Klingberg T, Jacobsen RB, Gabrieli JD (2000) A resource model of the neural basis of executive working memory. Proceedings of the National Academy of Sciences of the United States of America 97:3573–3578.

Cantlon JF, Brannon EM, Carter EJ, Pelphrey KA (2006) Functional imaging of numerical processing in adults and 4-y-old children. PLoS Biol 4:e125.

Chan MY, Park DC, Savalia NK, Petersen SE, Wig GS (2014) Decreased segregation of brain systems across the healthy adult lifespan. Proceedings of the National Academy of Sciences of the United States of America 111:E4997–5006.

Christodoulou C, DeLuca J, Ricker JH, Madigan NK, Bly BM, Lange G, Kalnin AJ, Liu WC, Steffener J, Diamond BJ, Ni AC (2001) Functional magnetic resonance imaging of working memory impairment after traumatic brain injury. J Neurol Neurosurg Psychiatry 71:161–168.

Cocchi L, Halford GS, Zalesky A, Harding IH, Ramm BJ, Cutmore T, Shum DH, Mattingley JB (2014) Complexity in relational processing predicts changes in functional brain network dynamics. Cereb Cortex 24:2283–2296.

Cohen JR, D’Esposito M (2016) The Segregation and Integration of Distinct Brain Networks and Their Relationship to Cognition. J Neurosci 36:12083–12094.

Cohen JR, Gallen CL, Jacobs EG, Lee TG, D’Esposito M (2014) Quantifying the reconfiguration of intrinsic networks during working memory. PloS one 9:e106636.

Cole MW, Reynolds JR, Power JD, Repovs G, Anticevic A, Braver TS (2013) Multi-task connectivity reveals flexible hubs for adaptive task control. Nat Neurosci 16:1348–1355.

D’Esposito M, Postle BR, Ballard D, Lease J (1999a) Maintenance versus manipulation of information held in working memory: An Event-Related fMRI study. Brain and cognition 41:66–86.

D’Esposito M, Zarahn E, Aguirre GK, Rypma B (1999b) The effect of normal aging on the coupling of neural activity to the bold hemodynamic response. NeuroImage 10:6–14.

Davis SW, Luber B, Murphy DLK, Lisanby SH, Cabeza R (2017) Frequency-specific neuromodulation of local and distant connectivity in aging and episodic memory function. Hum Brain Mapp.

De Pisapia N, Braver TS (2008) Preparation for integration: the role of anterior prefrontal cortex in working memory. Neuroreport 19:15–19.

Eldreth DA, Patterson MD, Porcelli AJ, Biswal BB, Rebbechi D, Rypma B (2006) Evidence for multiple manipulation processes in prefrontal cortex. Brain Res 1123:145–156.

Eriksson J, Vogel EK, Lansner A, Bergstrom F, Nyberg L (2015) Neurocognitive Architecture of Working Memory. Neuron 88:33–46.

Fornito A, Harrison BJ, Zalesky A, Simons JS (2012) Competitive and cooperative dynamics of large-scale brain functional networks supporting recollection. Proceedings of the National Academy of Sciences of the United States of America 109:12788–12793.

Golde M, von Cramon DY, Schubotz RI (2010) Differential role of anterior prefrontal and premotor cortex in the processing of relational information. NeuroImage 49:2890–2900.

Gruber O, Indefrey P, Steinmetz H, Kleinschmidt A (2001) Dissociating neural correlates of cognitive components in mental calculation. Cerebral cortex 11:350–359.

Han SH, Kim MS (2004) Visual search does not remain efficient when executive working memory is working. Psychological science 15:623–628.

Harvey BM, Klein BP, Petridou N, Dumoulin SO (2013) Topographic representation of numerosity in the human parietal cortex. Science 341:1123–1126.

Hearne LJ, Cocchi L, Zalesky A, Mattingley JB (2017) Reconfiguration of Brain Network Architectures between Resting-State and Complexity-Dependent Cognitive Reasoning. The Journal of neuroscience: the official journal of the Society for Neuroscience 37:8399–8411.

Hullett PW, Hamilton LS, Mesgarani N, Schreiner CE, Chang EF (2016) Human Superior Temporal Gyrus Organization of Spectrotemporal Modulation Tuning Derived from Speech Stimuli. J Neurosci 36:2014–2026.

Knuth DE (1976) Mathematics and computer science: coping with finiteness. Science 194:1235–1242.

Kraha A, Turner H, Nimon K, Zientek LR, Henson RK (2012) Tools to support interpreting multiple regression in the face of multicollinearity. Frontiers in psychology 3:44.

Libby LA, Hannula DE, Ranganath C (2014) Medial temporal lobe coding of item and spatial information during relational binding in working memory. J Neurosci 34:14233–14242.

Mumford JA, Poline JB, Poldrack RA (2015) Orthogonalization of regressors in FMRI models. PloS one 10:e0126255.

Overath T, McDermott JH, Zarate JM, Poeppel D (2015) The cortical analysis of speech-specific temporal structure revealed by responses to sound quilts. Nat Neurosci 18:903–911.

Park J, Li R, Brannon EM (2014) Neural connectivity patterns underlying symbolic number processing indicate mathematical achievement in children. Dev Sci 17:187–202.

Piazza M, Pinel P, Le Bihan D, Dehaene S (2007) A magnitude code common to numerosities and number symbols in human intraparietal cortex. Neuron 53:293–305.

Postle BR, Ferrarelli F, Hamidi M, Feredoes E, Massimini M, Peterson M, Alexander A, Tononi G (2006) Repetitive transcranial magnetic stimulation dissociates working memory manipulation from retention functions in the prefrontal, but not posterior parietal, cortex. Journal of cognitive neuroscience 18:1712–1722.

Ranganath C, DeGutis J, D’Esposito M (2004a) Category-specific modulation of inferior temporal activity during working memory encoding and maintenance. Brain research Cognitive brain research 20:37–45.

Ranganath C, Cohen MX, Dam C, D’Esposito M (2004b) Inferior temporal, prefrontal, and hippocampal contributions to visual working memory maintenance and associative memory retrieval. J Neurosci 24:3917–3925.

Rubinov M, Sporns O (2010) Complex network measures of brain connectivity: uses and interpretations. NeuroImage 52:1059–1069.

Rypma B (2006) Factors controlling neural activity during delayed-response task performance: testing a memory organization hypothesis of prefrontal function. Neuroscience 139:223–235.

Rypma B, D’Esposito M (1999) The roles of prefrontal brain regions in components of working memory: effects of memory load and individual differences. Proceedings of the National Academy of Sciences of the United States of America 96:6558–6563.

Rypma B, Prabhakaran V, Desmond JE, Glover GH, Gabrieli JD (1999) Load-dependent roles of frontal brain regions in the maintenance of working memory. NeuroImage 9:216–226.

Schedlbauer AM, Copara MS, Watrous AJ, Ekstrom AD (2014) Multiple interacting brain areas underlie successful spatiotemporal memory retrieval in humans. Sci Rep 4:6431.

Simony E, Honey CJ, Chen J, Lositsky O, Yeshurun Y, Wiesel A, Hasson U (2016) Dynamic reconfiguration of the default mode network during narrative comprehension. Nat Commun 7:12141.

Stevens AA, Tappon SC, Garg A, Fair DA (2012) Functional brain network modularity captures inter- and intra-individual variation in working memory capacity. PloS one 7:e30468.

Tournier JD, Calamante F, Connelly A (2007) Robust determination of the fibre orientation distribution in diffusion MRI: non-negativity constrained super-resolved spherical deconvolution. NeuroImage 35:1459–1472.

Tournier JD, Calamante F, Gadian DG, Connelly A (2004) Direct estimation of the fiber orientation density function from diffusion-weighted MRI data using spherical deconvolution. NeuroImage 23:1176–1185.

Tzourio-Mazoyer N, Landeau B, Papathanassiou D, Crivello F, Etard O, Delcroix N, Mazoyer B, Joliot M (2002) Automated anatomical labeling of activations in SPM using a macroscopic anatomical parcellation of the MNI MRI single-subject brain. NeuroImage 15:273–289.

Wylie GR, Javitt DC, Foxe JJ (2004) Don’t think of a white bear: an fMRI investigation of the effects of sequential instructional sets on cortical activity in a task-switching paradigm. Hum Brain Mapp 21:279–297.

Yeh CH, Smith RE, Liang X, Calamante F, Connelly A (2016) Correction for diffusion MRI fibre tracking biases: The consequences for structural connectomic metrics. NeuroImage 142:150–162.

